# Alterations in sialic-acid *O*-acetylation glycoforms during murine erythrocyte development

**DOI:** 10.1101/469254

**Authors:** Vinay S. Mahajan, Faisal Alsufyani, Hamid Mattoo, Ian Rosenberg, Shiv Pillai

## Abstract

9-*O*-acetylation of sialic acid is a common modification that plays important roles in host-pathogen interactions. CASD1 has been described as a sialate-*O*-acetyltransferase and has been shown to be essential for 9-*O*-acetylation of sialic acid in some cell lines *in vitro*. In this study, we used knockout mice to confirm that CASD1 is indeed responsible for 9-*O*-acetylation of sialic acids *in vivo*. We observed a complete loss of 9-*O*-acetylation of sialic acids on the surface of myeloid, erythroid and CD4^+^ T cells in *Casd1*-deficient mice. Although 9-*O*-acetylation of sialic acids on multiple hematopoietic lineages was lost, there were no obvious defects in hematopoiesis. Interestingly, red blood cells from *Casd1*-deficient mice also lost reactivity to TER-119, a rat monoclonal antibody that is widely used to mark the murine erythroid lineage. The sialic acid glyco-epitope recognized by TER-119 on red blood cells was sensitive to the sialic acid *O*-acetyl esterase activity of the hemagglutinin esterase from bovine coronavirus but not to the corresponding enzyme from the influenza C virus. During erythrocyte development TER-119^+^ Ery-A and Ery-B cells could be stained by catalytically inactive bovine coronavirus hemaggutinin-esterase but not by the inactive influenza C hemagglutinin esterase, while TER-119^+^ Ery-C stage cells and mature erythrocytes were recognized by both virolectins. These results suggest that throughout murine erythrocyte development, cells of the erythroid lineage express a glycoconjugate bearing a modified 7,9-di-*O*-acetyl form of sialic acid, that is recognized specifically by the bovine coronavirus lectin and not by the influenza C hemagglutinin, and this modified sialic acid moiety is a component of the TER-119 epitope. As erythrocytes mature, the surface of Ery-C cells and mature erythrocytes also acquires a distinct CASD1-dependent 9-*O*-acetyl sialic acid moiety that can be recognized by virolectins from both influenza C and bovine coronavirus that are specific for 9-*O*-acetyl sialic acid.

## INTRODUCTION

Sialic acid, a monosaccharide with a nine-carbon backbone, is widely found in nature as a terminal sugar of animal and microbial glycoproteins and glycolipids. It plays important roles in animal development, cellular recognition processes and host-pathogen interactions (Schauer, 2009). A common modification of sialic acid is 9-*O*-acetylation, which has been implicated in sialoglycan recognition and ganglioside biology. The hemagglutinins or hemagglutinin-esterases of type C influenza viruses and certain nidoviruses specifically recognize the 9-*O*-acetylated form of sialic acid, and this form of sialic acid is required for the entry of these viruses into human cells. Although we had previously inferred a role for sialic acid 9-*O*-acetylation in regulating Siglec-2 (Cariappa et al., 2009), we have demonstrated that many of the phenotypes on the basis of which earlier conclusions were made, actually resulted from a mutation that was traceable to a commercially available C57BL/6 strain of mice (Mahajan et al., 2016).

Previous studies had shown that a biochemical sialate-*O*-acetyl transferase activity exists in Golgi preparations. Using knock-down and over-expression studies in a human cell line, Arming et al. showed that CASD1 contributed to the 7-*O*-acetylation of GD3 (Arming et al., 2010). Baumann et al. inactivated *Casd1* in the near-haploid myeloid cell line HAP1 using CRISPR/Cas9 based genome editing and demonstrated the loss of 9-*O*-acetyl moieties from sialo-glyconjugates in HAP1 cells lacking CASD1 (Baumann et al., 2015). Their studies implicated CASD1 in directly transferring acetyl moieties from acetyl-CoA to CMP-sialic acid. Whether CMP-sialic acid is the preferred substrate for the O-acetylation of sialic acid *in vivo* however remains to be established. In this report, we describe a mouse strain in which *Casd1* has been deleted in the germline. Analysis of these mice demonstrated the loss of 9-*O*-acetylation of sialic acid on the surface of all hematopoietic cells examined. Based on the differential specificity of hemagglutinin esterases from Influenza C and bovine coronavirus, we demonstrate that the murine erythroid lineage-specific epitope recognized by the TER-119 monoclonal antibody is comprised of a 7,9-di-*O*-acetyl sialic acid glycoform that is completely lost in *Casd1*-deficient mice. The 7,9-di-*O*-acetyl sialic acid glycoform can be detected early in erythroid differentiation from the Ery-A and Ery-B stages onward. We also observed that a distinct 9-*O*-acetyl sialic acid moiety also decorates cells of the erythroid lineage from the Ery C stage onward including in mature red blood cells.

## RESULTS

### Targeted deletion of *Casd1* in mice results in a loss of 9-*O*-acetyl sialic acid in mononuclear hematopoietic cells

Prior *in vitro* studies have suggested that CASD1 is a sialic acid 9-*O*-acetyl transferase. To assess the function of 9-*O*-acetyl sialic acids *in vivo*, we generated *Casd1*-deficient mice (**Fig 1A**). The influenza C hemagglutinin-esterase (CHE) that is known to be specific for 9-*O*-acetyl sialic acid was expressed as an Fc-fusion protein (CHE-Fc) (Muchmore and Varki, 1987). Inactivation of the esterase domain of CHE-Fc with diisopropyl fluorophosphate (DFP) yields CHE-FcD which has been widely used as a specific probe of 9-*O*-acetyl sialic acid (Klein et al., 1994). Using the CHE-FcD reagent, we evaluated the effect of *Casd1* deficiency on sialic acid 9-*O*-acetylation in all hematopoietic lineages, focusing our analysis on the spleen, bone marrow and lymph nodes.

**Figure 1:**
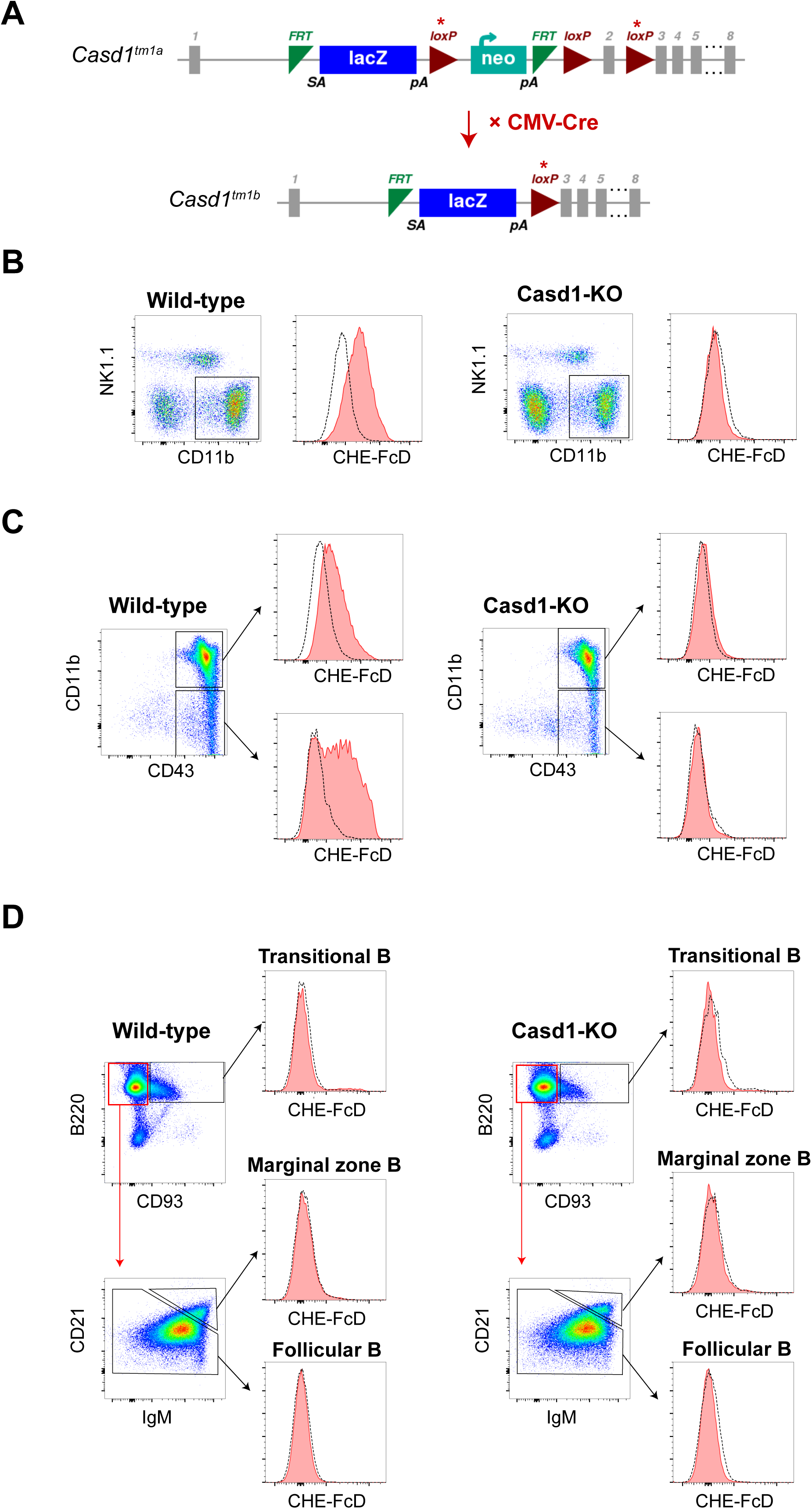
Loss of 9-*O*-acetyl sialic acid in myeloid cells in *Casd1*-deficient mice. (A) Generation of *Casd1* knockout mice. (B) Flow cytometry analysis of 9-*O*-acetylation CD11b^+^ (TER-119^−^ B220^−^ Thy1.2^−^ NK1.1^−^) myeloid cells using CHE-FcD probe in spleens of *Casd1*-deficient and wild-type (C57BL/6J) mice. Dotted lines depict control staining with secondary antibody only. (C) 9-O acetylation of CD43^+^ CD11b^+^ cells (TER-119^−^ CD71^−^ B220^−^ Thy1.2^−^) by CHE-FcD staining in bone marrows of *Casd1*-deficient and wild-type (C57BL/6J) mice. (D) CHE-FcD staining on B220^+^ CD93^+^ transitional B cells, B220^+^ CD19^+^ CD21^lo^ IgM^+^ follicular B cells and CD21^hi^ IgM^hi^ marginal zone B cells in the spleens of wild-type (C57BL/6J) or *Casd1*-deficient mice.

Wild-type myeloid cells expressing CD11b exhibit robust levels of 9-*O*-acetyl sialic acid, but this carbohydrate modification was completely lost in splenic CD11b^+^ myeloid cells from *Casd1*-deficient mice (**Fig 1B**). CD11b^+^ cells in the bone marrow also express high levels of the mucin-like glycoprotein CD43, suggesting that much of the 9-*O*-acetylated sialic acid found on the surface of murine myeloid cells may decorate this glycoprotein. Indeed, mucin-like glycoproteins such as CD43 and CD45RB have been shown to bear 9-*O*-acetylated sialic acids on mononuclear cells in the spleen (Krishna and Varki, 1997). No detectable 9-*O*-acetyl sialic acid was found on the CD11b^+^CD43^+^ population in the bone marrows of *Casd1*-deficient mice (**Fig 1C**). A large proportion of CD43^+^ CD11b^−^ non-myeloid cells, largely hematopoietic progenitors in the bone marrow, also express surface 9-*O*-acetyl sialic acid in wild type mice but this was completely lost in *Casd1*-deficient mice (**Fig 1C**).

We found no detectable levels of 9-*O*-acetyl sialic acid on B220^+^ CD93^+^ transitional B cells, B220^+^CD19^+^CD21^lo^ IgM^+^ follicular B cells and CD21^hi^ IgM^hi^ marginal zone B cells in the spleen of wild-type (C57BL/6J) or *Casd1*-deficient (**Fig 1D**). We also analyzed B220^+^ B cells in the bone marrow and lymph nodes, and found no detectable levels of 9-*O*-acetyl sialic acid on these cells in wild-type or *Casd1*-deficient mice (**Supplementary Figs 1A & 1B**).

The CHE-FcD probe revealed no appreciable levels of 9-*O*-acetyl sialic acids on CD8^+^ T cells or NK cells (**Figs 2A & 2B**) in the spleens of wild-type or *Casd1*-deficient mice. Similar results were observed in the lymph nodes (**Supplementary Figs 2A & 2B**). CD4^+^ T cells express high levels of 9-*O*-acetylated sialic acids (**Fig 2A**), which is partly on mucin-type glycoproteins and partly on gangliosides (Krishna and Varki, 1997). We found that naive CD44^lo^CD62L^hi^ CD4^+^ T cells had higher levels of 9-*O*-acetyl sialic acid than CD44^hi^CD62L'° effector memory CD4^+^ T cells (**Fig 2C**). The 9-*O*-acetylation of sialic acid normally observed on both naive and memory wild-type CD4^+^ T cells in the spleen was completely lost in *Casd1*-deficient mice (**Figs 2A & 2C**). The 9-*O*-acetyl sialic acid on wild-type CD4^+^ T cells was also eliminated by influenza C hemagglutinin esterase (CHE-Fc) treatment (**Fig 2D**).

**Figure 2:**
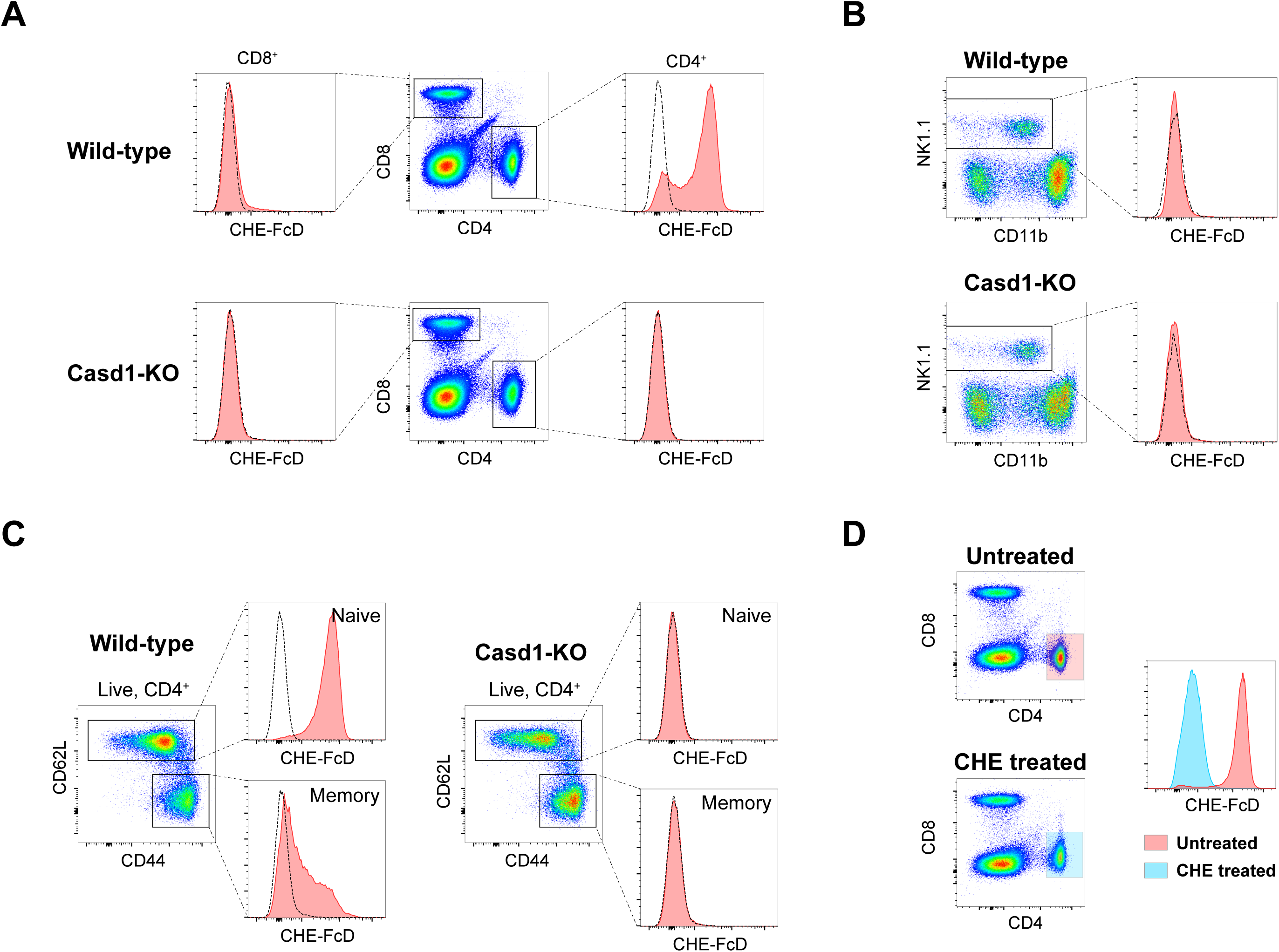
Loss of 9-*O*-acetylation in CD4^+^ T cells in *Casd1*-deficient mice. (A) Analysis of 9-*O*-acetylation by CHE-FcD staining of CD4^+^ and CD8^+^ T cells in spleens of *Casd1*-deficient and wild-type (C57BL/6J) mice using flow cytometry. Dotted lines depict control staining with secondary antibody only. (B) CHE-FcD staining on NK1.1^+^ (TER-119^−^ B220^−^ Thy1.2^−^) cells from the spleens of wild-type (C57BL/6J) or *Casd1*-deficient mice. (C) CHE-FcD staining of naive CD62L^+^ CD4^+^ T cells and memory CD44^+^ CD4^+^ T cells for 9-*O*-acetylation in spleens of *Casd1*-deficient and wild-type (C57BL/6J) mice. (D) CD4^+^ T cells from the spleen before and after treatment by Influenza C hemagglutinin esterase (CHE-Fc) at 37°C for 1 hour then stained by CHE-FcD for 9-*O*-Acetylation.

### The TER-119 monoclonal antibody fails to bind to *Casd1*-deficient murine erythrocytes and erythrocyte progenitors

Erythrocytes from *Casd1*-deficient mice also lost reactivity to CHE-FcD (**Fig 3A**). Despite the loss of 9-*O*-acetylation on erythrocytes, there was no evidence of anemia or other obvious change in RBC physiology. Surprisingly, *Casd1*-deficient erythrocytes, which lack 9-*O*-acetyl sialic acid based on the lack of binding to CHE-FcD, also lost reactivity to TER-119, suggesting that this antibody may recognize a 9-*O*-acetyl sialic acid glyco-epitope (**Fig 3A**). A previous study on the identification of the target of TER-119 suggested that this antibody can immunoprecipitate proteins of 110 kDa, 60 kDa, 52 kDa and 32 kDa from the erythrocyte membrane (Kina et al., 2000). Of these, only the 52 kDa protein was detectable in a Western Blot using TER-119. However, the identity of this protein was not deciphered, and it was suspected to be a glycophorin A associated molecule.

**Figure 3:**
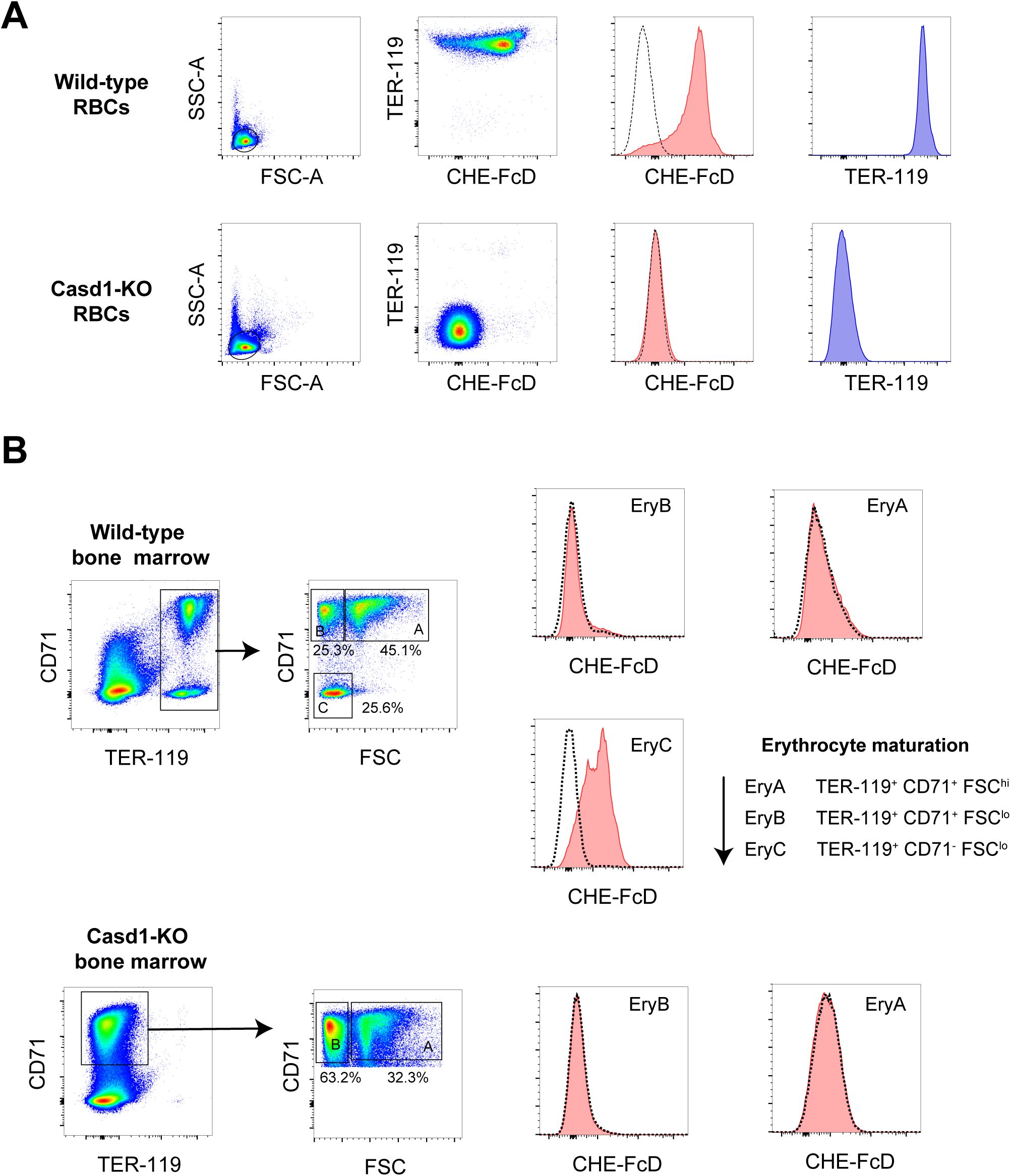
TER-119 monoclonal antibody fails to bind *Casd1*-deficient murine erythrocyte. (A) TER-119 and CHE-FcD staining on circulating red blood cells from *Casd1*-deficient and wild-type (C57BL/6J) mice. (B) Flow cytometric analysis of CHE-FcD levels on erythroid precursors in the bone marrow of wild-type (C57BL/6J) and *Casd1*-deficient mice. Please note that due to the loss of TER-119 in the erythroid lineage in *Casd1*-deficient mice, only the Ery-A and Ery-B precursors but not the Ery-C precursors can be definitively identified in *Casd1*-deficient mice using CD71 expression and FSC.

Erythrocyte precursors in the bone marrow can be identified based on CD71 and TER-119 staining and are divided into the Ery-A (TER-119^+^ CD71^+^ FSC^hi^), Ery-B (TER-119^+^ CD71^+^ FSC^lo^) and Ery-C (TER-119^+^ CD71^−^ FSC^lo^) flow cytometric fractions based on CD71, TER-119 and forward scatter (FSC) (**Fig 3B**) (Chen et al., 2009; Koulnis et al., 2011). We found that surface 9-*O*-acetyl sialic acid identified using CHE-FcD staining appears only at the most mature Ery-C stage in the erythroid lineage. We did not observe any CD71 positive cells that lacked TER-119 in the wild-type mouse bone marrow, suggesting that CD71 and FSC alone could be used to identify the Ery-A and Ery-B erythroblast fractions in the bone marrow of *Casd1*-deficient mice (that lack the TER-119 epitope) (**Fig 3B**). As in wild-type mice, in the bone marrow of *Casd1*-deficient mice, there was also no detectable surface 9-*O*-acetylation on CD71^+^ Ery-A and Ery-B erythroid progenitors based upon staining with CHE-FcD (**Fig 3B**). Our flow cytometry analysis suggests that *Casd1*-deficient mice exhibit a pan-hematopoietic loss of 9-*O*-acetyl sialic acids in the myeloid, lymphoid and erythroid lineages.

### The TER-119 epitope is sensitive to the hemagglutinin esterase from bovine coronavirus but not influenza C

Given that the *Casd1*-deficient mice exhibited a complete loss of TER-119 staining in the erythroid lineage, we considered the possibility that the TER-119 monoclonal antibody recognized a 9-*O*-acetyl sialic acid dependent epitope. However, the Ery-A and Ery-B erythroid precursors that normally express the TER-119 epitope in wild-type mice do not express any 9-*O*-acetyl sialic acid that is detectable by CHE-FcD staining. Therefore, we decided to examine the relationship between 9-*O*-acetyl sialic acid and the TER-119 epitope in greater detail. The hemagglutinin-esterase from bovine coronavirus Mebus strain (BHE) also recognizes 9-*O*-acetyl sialic acid, but in a manner that is structurally distinct from CHE (Zeng et al., 2008). Furthermore, unlike CHE, the BHE lectin recognizes 7,9-di-*O*-acetyl sialic acid in addition to 9-*O*-acetyl sialic acid (Langereis et al., 2015; Bakkers et al., 2016). Therefore, we expressed BHE as an Fc-fusion protein (BHE-Fc) analogous to the approach used to generate CHE-Fc. In addition, we also generated a catalytically inactive form of BHE-Fc bearing the S40A substitution (BHE-Fc-S40A) whose ability to bind 9-*O*-acetyl sialic acids is unhindered as previously described (Zeng et al., 2008). We found that the TER-119^+^ CD71^+^ Ery-A and Ery-B precursors from wild-type mice exhibited strong staining with BHE-Fc-S40A but not with CHE-FcD (**Fig 3B and 4A**). This suggested to us that the TER-119 epitope may be recognized by the BHE virolectin but not by CHE. We next used murine RBCs to examine the interactions of the TER-119 epitope with CHE and BHE.

**Figure 4:**
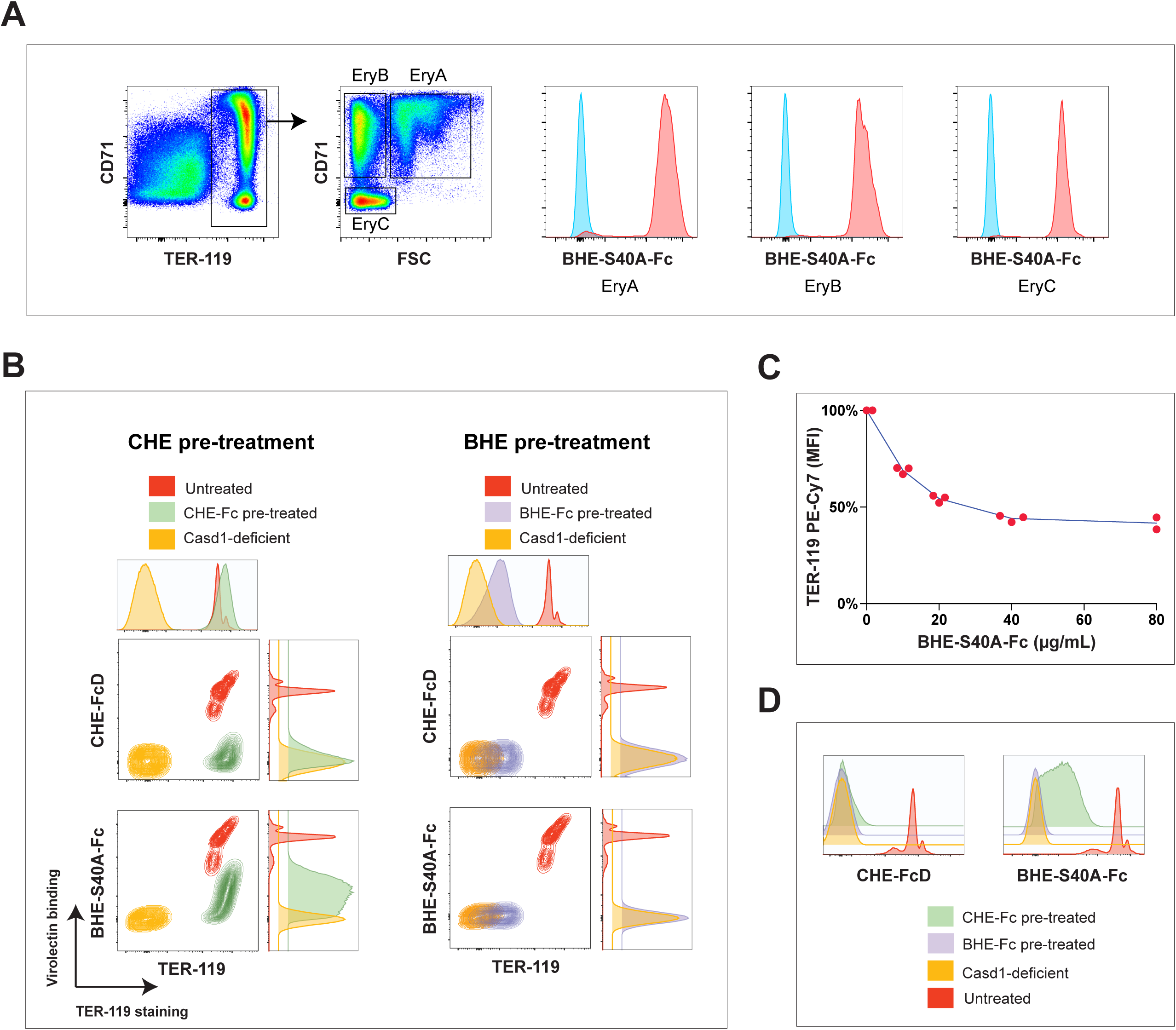
TER-119 epitope is sensitive to the hemagglutinin esterase from bovine coronavirus (BHE) but not influenza C virus (CHE) (A) BHE-S40A-Fc staining on Ery-A, Ery-B and Ery-C precursors from wild-type (C57BL/6J) mice. (B) Analysis of the binding to TER-119 and virolectins from bovine coronavirus (BHE-S40A-Fc) and influenza C virus (CHE-FcD) to wild-type and *Casd1*-deficient red blood cells with and without treatment with two respective viral esterases (CHE and BHE). (C) Competitive binding of the fluorescently labeled TER-119 antibody to red blood cells in the presence of increasing concentrations of the enzymatically inactive bovine coronavirus virolectin (BHE-S40A-Fc). Red blood cells were pre-incubated in the presence of the catalytically inactive virolectin for one hour at 37°C and subsequently exposed to 2 μg/mL of labeled TER-119 antibody for 10 minutes in the continued presence of the virolectin. (D) CHE-FcD or BHE-S40A-Fc virolectin staining before and after CHE of BHE treatment of red blood cells from wild-type or *Casd1*-deficient mice.

Influenza C hemagglutinin esterase (CHE-Fc) pre-treatment eliminated 9-*O*-acetyl sialic acid-dependent CHE-FcD binding on murine erythrocytes rendering them indistinguishable from *Casd1*-deficient erythrocytes based on CHE-FcD staining (**Fig 4B**). Yet, the TER-119 epitope on murine erythrocytes was surprisingly resistant to treatment with CHE (**Fig 4B**). As a control, and as previously shown by Varki et al., we also showed that CHE-Fc pre-treatment completely eliminated CHE-FcD binding on CD4^+^ T lymphocytes (**Fig 2D**). Interestingly, the TER-119 epitope on murine RBCs was sensitive to BHE-Fc but not CHE-Fc esterase treatment (**Fig 4B**). In contrast, pretreatment with an enzymatically inactive version of BHE-Fc bearing the S40A catalytic site mutation (BHE-S40A-Fc) had a negligible effect on TER-119 staining (data not shown).

However, in the setting of a competitive binding assay, the lectin domain of BHE modestly hampered TER-119 binding to its epitope in a dose-dependent manner (**Fig 4C**). Given that both CHE and BHE have evolved as receptor-destroying enzymes that recognize 9-*O*-acetyl sialic acids, it was not surprising that both CHE-Fc and BHE-Fc pre-treatment of mouse RBCs was able to eliminate the corresponding virolectin receptor binding (**Fig 4D**). Although treatment with BHE-Fc completely eliminated CHE-FcD binding, CHE only partially eliminated BHE-S40A-Fc binding (**Fig 4D**). These data are consistent with the notion that the BHE virolectin and enzyme has the ability to recognize modified 9-*O*-acetyl sialic acid ligands that are not recognized by CHE. Since BHE has been shown to bind to 7,9-di-*O*-acetyl sialic acid while CHE cannot (Langereis et al., 2015; Bakkers et al., 2016), our results support the notion that 7,9 di-*O*-acetyl sialic acid is a component of the TER-119 epitope and is expressed in the erythroid lineage from the Ery-A stage onwards.

## DISCUSSION

Prior studies of Baumann *et al* have shown that CASD1 catalyzes the transfer of acetyl moieties from acetyl-CoA to CMP-sialic acid through the formation of a covalent acetyl-enzyme intermediate (Baumann et al., 2015). Of note, CASD1 is unable to use the unactivated form of sialic acid or other sialic acid that is already incorporated into sialoconjugates (e.g. fetuin (N-linked), bovine submandibular mucin (O-linked), 2,3/2,6-sialolactose, or GD3) as substrates. The authors proposed that the 9-*O*-acetylated CMP-sialic acid is subsequently incorporated into sialoglycans. CASD1 is widely expressed in most tissues (based on the Protein and RNA Expression Atlas); this may indicate that most 9-*O*-acetyl sialoglycans are generated in a CASD1-dependent manner (Uhlen et al., 2015). Salvage of sialic acids from dietary sialogycans may serve as an additional source of sialic acid, but it remains unclear if the 9-*O*-acetyl moiety is preserved during this process. Our results suggest that CASD1 is necessary for the synthesis of 9-*O*-acetyl sialic acid at least in all hematopoietic cells.

In mammals, O-acetylation of sialic acid has been observed at the 4, 7 and 9 positions. In addition enzymatic activity corresponding to transferases and esterases capable of adding or removing these substitutions have been successfully enriched from subcellular fractions of tissues rich in O-acetylated sialic acids. Among these, CASD1 is the only mammalian enzyme that has been cloned and biochemically characterized; we have described the generation of a *Casd1* knockout mouse in this study. However, our knowledge of the mammalian enzymes O-acetylating sialic acids remains incomplete and there has been a debate about the number of enzymes involved in generating the pool of 9-*O*-acetyl sialic acids *in vivo*. It remains to be established whether incorporation of 9-*O*-acetylated form of CMP-NANA into glycans is the only mechanism of generating 9-*O*-acetylated sialogycans. The complete loss of 9-*O*-acetyl sialic acids on the surface of erythroid, myeloid and CD4 T lineages suggests that at least in unmanipulated *Casd1*-deficient mice, CASD1 may be the only enzyme capable of acetylating sialic acids in the 9-OH position on these cell types. Our results also suggest that CASD1 is required for the generation of 7,9-di-*O*-acetyl sialic acids in the murine erythroid lineage.

Based on our analysis of the differential binding or inactivation of the TER-119 epitope by BHE and CHE virolectins and receptor-destroying esterases, the TER-119 antibody appears to recognize an epitope bearing 7,9-di-*O*-acetyl sialic acid. The analysis of the ligand-bound structures of BHE and CHE shows that although there is some structural homology between the two hemagglutinins, they bind the 9-*O*-acetyl sialic acid ligand in opposite orientations (Zeng et al., 2008). Thus, it is theoretically possible that the conformation of the specific 9-*O*-acetyl sialic acid glycoform recognized by TER-119 is sterically constrained in the context of a particular glycan that favors binding and recognition by BHE over CHE. Although both BHE and CHE are lectins fused to receptor-destroying enzymes, it is particularly interesting that the esterase and hemagglutinin activities appear to have evolutionarily converged at the level of the specific glyco-conformations of 9-*O*-acetyl sialic acids. Thus study reveals an additional layer of complexity and diversity among the O-acetylated sialic acids and establishes the developmental regulation of the sequential acquisition of two different forms of 9-*O*-acetyl sialic acid during murine erythroid differentiation.

## METHODS

### Generation of Casd1 knockout mice

Mouse embryonic stem cells (C57/BL6N background) carrying a knockout-first targeted allele of *Casd1* (*Casd1^tm1a(EUCOMM)Hmgu^*) were obtained from the EUCOMM consortium. *Casd1* knock-in mice were generated at the BWH Transgenic Core facility. The resulting chimeric mice were crossed into C57BL/6NTac (Taconic) and interbred to homozygosity. They were then crossed into (B6.C-Tg(CMV-cre)1Cgn/J mice, (stock # 006054, Jackson Laboratory) to obtain the *Casd1^tm1b^* null allele, which lacks exon 2 of *Casd1*. Excision of the neomycin expression cassette (**Fig 1A**) was confirmed by PCR. The following primers were used for genotyping. The wild-type allele was specifically amplified using TCCTCCCTCACTGTTCCTTC and AGGTGGGGAGGAAAGACAGT (291bp) and the *Casd1^tm1b^* allele could be selectively amplified using TCCTCCCTCACTGTTCCTTC and GACAGCCAGTCACACAGCTT (5540bp). This allele was interbred to homozygosity to yield *Casd1^tm1b/tm1b^* mice (referred to as *Casd1*-KO mice in the accompanying figures). The *Casd1*-deficient mice were viable and fertile. The *Casd1* knockout colony was maintained by homozygous mating. Each experiment was performed at least twice on three mice each.

### Flow cytometry

Single cell suspensions were obtained from the blood, spleen, lymph nodes, thymus or bone marrow and flow cytometry analysis was performed using the following monoclonal antibodies: B220 (RA3-6B2), CD4 (GK1.5), CD8 (53-6.7), CD44 (IM7), CD62L (MEL-14), CD90.2 (30-H12), NK-1.1 (PK136), CD11b (M1/70), CD71 (RI7217), TER-119 (TER-119), CD11b (M1/70), CD3 (145-2C11), CD43 (S11), CD93 (AA4.1), CD19 (1D3), CD21 (7G6) and CD16/CD32 (2.4G2). Monoclonal antibody clones are indicated in parentheses. All antibodies were obtained from Biolegend Inc. unless otherwise indicated. One representative flow cytometry plot is shown for each condition.

### Studies with coronavirus and influenza C virolectins

Influenza C hemagglutinin esterase (CHE) was recombinantly expressed as an Fc-fusion protein and the esterase activity of CHE was inactivated with di-isofluoropropyl phosphate to generate the CHE-FcD reagent as previously described (Muchmore and Varki, 1987). Bovine coronavirus hemagglutinin esterase (Mebus strain, Uniprot accession number, P15776.1) was cloned into the pFUSE-mFc2a vector (Invivogen) into the EcoRI and BglII sites. A catalytically dead variant of BHE-Fc (BHE-S40A-Fc) was generated by site-directed mutagenesis. The Fc-fusion proteins were expressed in 293T cells and affinity purified using protein G. The CHE-FcD probe recognizes 9-*O*-acetyl sialic acids, and the BHE-S40A-Fc virolectin can bind both 9-*O*-acetyl and 7,9-di-*O*-acetyl sialic acids (Langereis et al., 2015; Bakkers et al., 2016). Briefly, a two-step staining procedure with CHE-FcD as the primary and anti-human IgG1-PE (clone HP6017) as the secondary or with BHE-S40A-Fc as the primary and anti-mouse IgG2a-FITC (clone RMG2a-62) as the secondary antibody were used.

To assess the effect of CHE or BHE on 9-*O*-acetyl sialic acids on murine erythrocytes, a 1:100 dilution of whole blood in PBS, was treated with 10μg/mL of CHE-Fc or BHE-Fc at 37°C for 1 hour. The cells were washed twice with PBS at room temperature and stained with CHE-FcD as described above. Analysis of competitive binding between TER-119 and BHE-S40A-Fc was performed as follows. A 1:100 dilution of whole blood in PBS was incubated with varying concentrations of BHE-S40A-Fc at 37°C for one hour followed by the addition of 2 μg/mL of labeled TER-119 antibody for 10 minutes in the continued presence of the virolectin. The cells were washed twice with ice cold PBS and analyzed by flow cytometry.

## ACKNOWLEDGEMENTS

This work was supported by research grants from the National Institutes of Health to SP (AI064930 and AI110495) and VM (AI113163).

